# Biotic and abiotic factors predict the biogeography of soil microbes in the Serengeti

**DOI:** 10.1101/2020.02.06.936625

**Authors:** Bo Maxwell Stevens, Derek Lee Sonderegger, Nancy Collins Johnson

**Author notes:** **Corresponding Author:** Bo Maxwell Stevens.

## Abstract

Field-based observational research is the first step in understanding the factors that predict the biogeography and community structure of soil microbes. The Serengeti National Park in Tanzania is an ideal location for this type of research because active volcanoes generate strong environmental gradients due to ash deposition and a rain shadow. Also, as one of the last remaining naturally grazed ecosystems on Earth, the Serengeti provides insights about the influence of herbivory on microbial communities. We used 16S rRNA amplicons to characterize bacterial and archaeal communities in soils from a 13-year herbivore removal experiment to study the influence of environmental factors and grazing on the natural distribution of soil microbes. We collected soil samples from seven sites, each with three naturally grazed plots and three plots that were fenced to prevent grazing by large mammalian herbivores. Soil fertility (phosphorus, nitrogen, iron, calcium, organic matter), texture, and pH were measured at each plot. Beta diversity of bacterial and archaeal communities was most strongly correlated with soil texture (R^2^ = 32.4%). The abundance of many operational taxonomic units (OTUs) were correlated with soil texture, and the evenness of taxa within samples increased with fine-textured soil. Removal of grazing shifted community structure, with 31 OTUs that were significant indicator taxa of the ungrazed treatment and three OTUs that were significant indicators of the grazed treatment.

**Importance:** Our results show that in this regional scale study, soil texture was the best environmental predictor, and grazing by large mammals also structures bacterial and archaeal communities. When large mammals are removed, as humans have been doing for millenia, there are cascading effects into the microbial world that can influence ecosystem functions like carbon and nitrogen cycles. These empirical findings from a natural tropical savannah can help inform models of the global distribution and function of soil microbes.

## Introduction

Soil bacteria and archaea serve critical functions in natural ecosystems. Many environmental factors can be used to predict the distribution and diversity of communities of bacteria within the soil (Fierer, 2017). Although recent studies have elucidated patterns in the structure of bacterial communities, disentangling their biogeography by determining the underlying mechanisms affecting the distribution of bacteria and archaea (hereafter referred to as microbes) remains a challenge. The Serengeti National Park in Tanzania provides an ideal environmental gradient for investigating the effects of abiotic and biotic factors on microbial communities. Four mechanisms underlying the biogeography of soil microbes have been identified: selection, dispersal, drift, mutation (Hanson *et al.*, 2012). The purpose of this study is to consider how environmental selection and dispersal by large animals influences the biogeography of soil microbes in the Serengeti. To accomplish this, we examined the diversity and distribution of microbial communities across a natural gradient of abiotic conditions that is superimposed on a replicated experiment that removed large mammalian herbivores for 13 years.

In the Serengeti, local topography and volcanic inputs have created inverse gradients of soil properties and precipitation (Anderson & Talbot, 1965; Ashley *et al.*, 2014). Active volcanoes to the east continue to influence the geologic context with eruptions as recent as 2008 (Sinclair & Arcese, 1995; Vaughan *et al.*, 2008). Ash deposits from the nearby Ngorongoro Volcanic Highlands create gradients of calcium, iron, phosphorus and pH (Anderson & Talbot, 1965; Ruess & Seagle, 1994; Ashley *et al.*, 2014). Many studies have found that pH is a significant driver of microbial community composition globally (Fierer & Jackson, 2006; Lauber *et al.*, 2009; Griffiths *et al.*, 2011; Kaiser *et al.*, 2016). Phosphorus availability is linked to soil pH, calcium, and iron because of chemical interactions on soil particles. Phosphorus availability is highest in neutral pH ranges. Calcium immobilizes phosphorus above neutral pH ranges (Schachtman *et al.*, 1998; Brady & Weil, 2017), and iron can immobilize phosphorus below a neutral pH, indirectly affecting the growth of microbes and plants (Vance *et al.*, 2003; Pigna & Violante, 2003). Increasing phosphorus availability can influence microbial communities by altering assemblages or increasing diversity (Beauregard *et al.*, 2009; Kuramae *et al.*, 2011, 2012; Tan *et al.*, 2012). Abiotic soil characteristics directly affect microbial communities by influencing resource availability and habitat structure, and they indirectly affect bacterial communities by influencing biotic factors such as vegetation and food webs.

Soil microbes exist within a complex web of organisms that includes large grazing animals with many potential mechanisms for affecting microbial communities. The Serengeti hosts one of the largest natural grazing ecosystems in the world with currently over two million large mammalian herbivores (Eby *et al.*, 2014). Moderate levels of grazing can alter soil nutrient concentrations by transporting nutrients and modifying plant communities (Anderson *et al*. 2007). In ecosystems with high soil fertility where nutrient cycling is dominated by bacteria, grazing may have a positive effect on decomposer bacteria (Bardgett & Wardle, 2003). In addition to the direct effects of urine and dung on soil biota (McNaughton *et al.*, 1997; Bardgett *et al.*, 1998), grazing can indirectly influence microbial diversity by stimulating net primary productivity and aboveground biomass (McNaughton, 1979, 1985), leading to increased photosynthesis and root exudation (Bardgett *et al.*, 1998). Also, grazing may influence the composition of plant communities which may alter the composition of microbial communities (Prober *et al.*, 2015). Comparison of soil microbes between grazed and ungrazed treatments in a long-term field experiment will help elucidate shifts in bacterial communities resulting from the cascade of biotic and abiotic responses to the activities of large mammalian herbivores.

Although no single factor consistently explains the biogeography of soil microbes, certain environmental variables tend to correspond with spatial patterns in the composition and abundance of bacteria. Soil organic matter, pH, redox status, soil moisture, nitrogen and phosphorus availability, and soil texture appear to be important predictors of the structure of soil microbial communities (Fierer, 2017). Our study tests the hypothesis that selection by many of these abiotic factors structures microbial communities. Also, we compared microbial communities inside and outside grazing exclosures to test the hypothesis that seasonal defoliation by large mammalian herbivores structure microbial communities. The goal of this study is to increase our understanding of the biogeography of soil microbes in a naturally grazed, tropical grassland. This knowledge can inform the development of general principles to help predict the structure and function of soil bacterial communities in changing environments.

## Methods

### Sites

In 1999, a herbivore removal experiment was installed at eight sites within the Serengeti National Park, Tanzania (Fig. 1) (Anderson *et al.*, 2007). Six plots (4 × 4 m) were established at each site with three randomly assigned plots open to grazing and the other three plots had 2 m tall chain-linked fences to exclude grazing by large animals including wildebeest, zebras, Thomson’s gazelles, buffalo, and topi (McNaughton, 1985; Anderson *et al.*, 2007). Theft of all the fences at one site in the eastern corridor prevented its inclusion in this study. Soil samples were collected from the remaining seven sites during the rainy season in May, 2012. Precipitation in the Serengeti National Park in 2012 was lowest in the southern (662 mm) and highest in the northern sites (1143 mm; Table 1).

**Figure 1.**
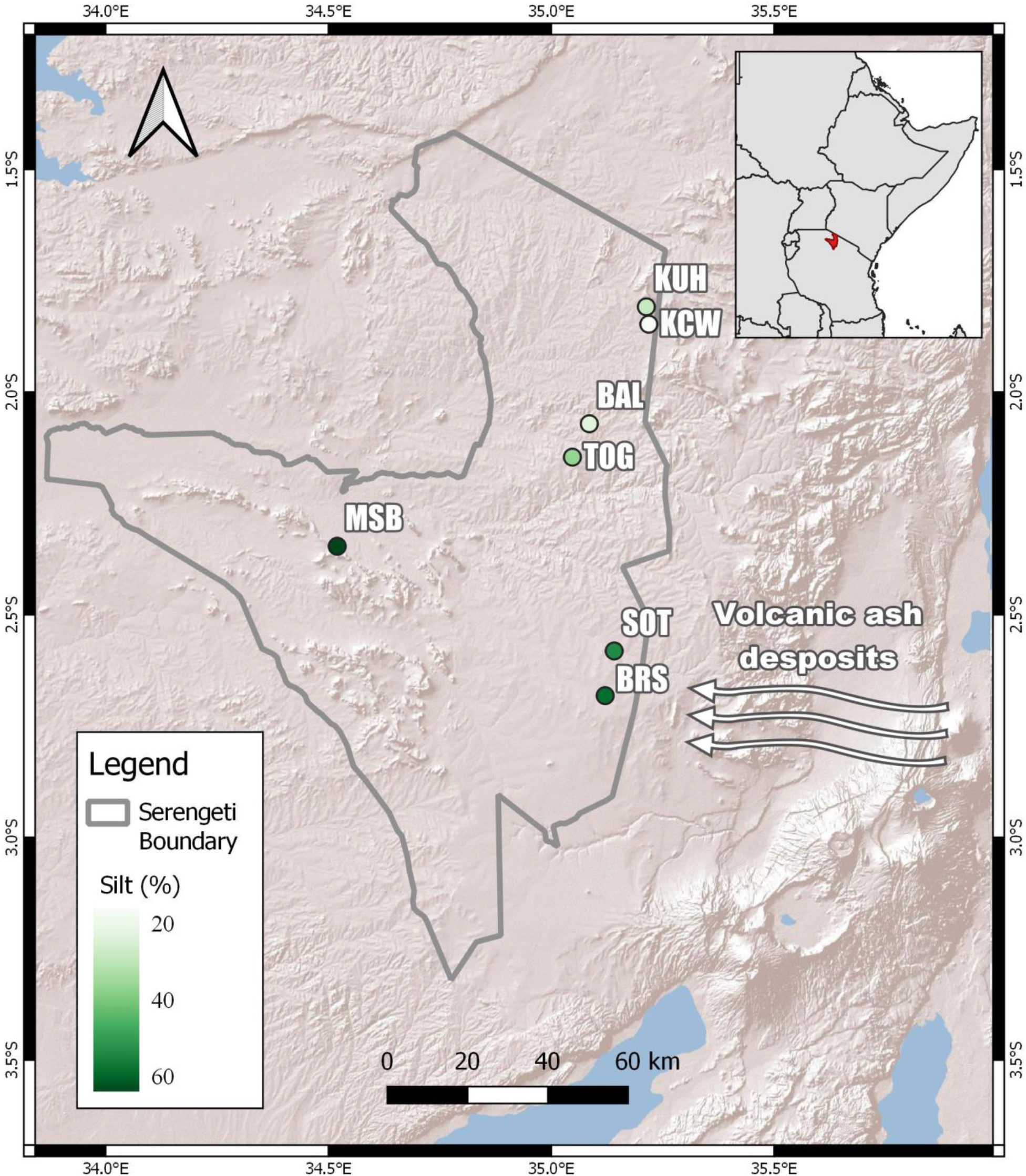
Locations of the seven study sites within the Serengeti National Park, Tanzania: Balatines (BAL), Barafu (BRS), Klein’s Camp West (KCW), Kuku Hills (KUH), Soit le Motonyi (SOT), Togora (TOG), and Musabi Plains (MSB). The gradient in soil texture is illustrated in green, darker color indicates higher percent silt.

### Soil analyses

Soil samples were collected from holes (approximately 15 cm deep) created by excavating grasses at all seven sites (n = 42). Within six hours of collection, soils were dried for 48 hours in a solar drier. After 2 weeks, the dry samples were brought to the laboratory and frozen for long-term storage. Frozen soil samples were dried at 103°C and sieved (< 2 mm). Soil organic matter was measured using loss on ignition, 2 g subsamples were weighed, heated to 550°C for 24 hours in a Lindberg HB muffle furnace (Lindberg/MPH, Riverside, MI 49084) then reweighed (Heiri *et al.*, 2001). Soil pH was measured potentiometrically in a 1:2.5 water:soil paste at the Soil Science Laboratory of Sokoine University of Agriculture in Morogoro, Tanzania (Klute, 1986). To measure total phosphorus, calcium, and iron concentrations, 0.3 g subsamples were ground and digested in 7 mL concentrated nitric acid and 3 mL 30% hydrogen peroxide in Milestone 900 Microwave Digestor (Ethos Inc., Bristol, United Kingdom). Samples were digested for 20 minutes and reached a maximum temperature of 425°C. Total soil phosphorus concentration converted to orthophosphate was quantified via colorimetry (Grimshaw, 1987) on a QuikChem 8000 Series FIA+ (Lachat Instruments, Milwaukee, WI 53218) using QuikChem Method 10-115-01-1-A. Total iron and calcium were measured on an AAnalyst 100 Atomic Absorption Spectrophotometer (Perkin Elmer, Waltham, MA 02451). Samples were compared to in-house standards and external standards produced by Ricca Chemical Company (Arlington, TX 76012) and Hach Company (Loveland, CO 80539).

Soil texture was determined using laser diffraction particle size analysis (Beuselinck *et al.*, 1998). Unsieved soil samples were suspended in water and analyzed on a LS 13 320 Series Laser Diffraction Particle Size Analyzer (Beckman Coulter Brea CA 92821). Particle sizes were grouped according USDA soil texture classification. Our soil textural classification figure was generated from R code available online (Hamilton, 2014). Soil bulk density and total soil nitrogen concentrations were obtained from previous analyses from the same plots (Antoninka *et al.*, 2015). Measurements of soil organic matter, total phosphorus, nitrogen, calcium, and iron were adjusted by bulk density.

### Molecular analysis

Amplicons were produced in a two-step protocol (Berry *et al.*, 2011). Samples were amplified in triplicate PCR reactions for the 16S v4 region using the universal bacterial primers 515F (5’-GTGCCAGCMGCCGCGGTAA-3’) and 806R (5’-GGACTACHVGGGTWTCTAAT-3’) (Bates *et al.*, 2011). First round reactions were performed in triplicate in 384 well plates. The 8 μL volumes contained the following: 1 μM each primer, 200 μM each dNTP (Phenix Research, Candler, NC), 0.01 U/μL Phusion HotStart II DNA Polymerase (Life Technologies), 1X HF Phusion Buffer (Life Technologies), 3 mM MgCl_2_, 6% glycerol, and 1 μL normalized template DNA. Cycling conditions were: 2 minutes at 95°C followed by 20 cycles of 30 seconds at 95°C, 30 seconds at 55°C, 4 minutes at 60°C. Triplicate reactions for each sample were pooled by combining 4 μL from each well, and 2 μL was used to check for results on an agarose gel. The remainder was diluted 10-fold and used as template in a second PCR reaction in which indexed tails (Caporaso *et al.*, 2012) were added. Second round reaction conditions were identical to the first round except only one reaction was conducted per sample and only 15 total cycles were performed. Indexed PCR products were purified using carboxylated magnetic beads (Rohland & Reich, 2012), quantified by PicoGreen fluorescence, and an equal mass of each sample was combined into a final sample pool. The pool was purified and concentrated, and subsequently quantified by quantitative PCR against Illumina DNA Standards (Kapa Biosystems, Wilmington, MA). Sequencing was carried out on a MiSeq Desktop Sequencer (Illumina Inc, San Diego, CA) running in paired end 2×150 mode. Upon acceptance, sequence will be archived in the NCBI Sequence Read Archive.

### Data analysis

The forward reads of the 16S amplicons were imported into QIIME 2 version 2018.11 (Bolyen *et al.*, 2018). Demultiplexing was carried out using minimum quality threshold of q20 and default parameters in QIIME 2. Based on quality threshold, visualized with FastQC version 11.7, reads were trimmed to 139 bases (Andrews, 2010). To determine phylogenetic diversity metrics, a rooted phylogenetic tree was created with MAFFT sequence alignment and FastTree in QIIME 2. QIIME 1.9.1 was used to filter samples below 0.005% abundance (Caporaso *et al.*, 2010; Bokulich *et al.*, 2013). To remove singletons by sample, the `otu_picking_workflow.sh` command in akutils v1.1.1 was performed (Andrews, 2018). Alpha and beta diversity metrics were performed with the q2-diversity plugin for QIIME 2 using the core-metrics-phylogenetic command with a sampling depth of 45000. We used Shannon diversity index to capture bacterial richness and evenness. To estimate alpha diversity with phylogenetic structure, we used Faith’s Phylogenetic Diversity index (Faith, 1992). To separate community evenness, we used Pielou’s evenness index, where values are constrained to 0 and 1, with higher values representing even abundance of community members. To generate figures, we used the scikit-bio 0.2.3 (http://scikit-bio.org), matplotlib 3.1.0, and seaborn 0.9.0 python packages. Upon acceptance, our environmental data, correlation matrix, and OTU table will be made publically available on Dryad Digital Repository (https://datadryad.org).

### Statistical analyses

In an attempt to account for spatial autocorrelation, beta diversity of bacterial communities were analyzed with single regressions (Anderson *et al.*, 2011) of Bayesian general linear mixed effects models using the “rstanarm” R package version 2.17.4 (Stan Development Team, 2017). Weighted and unweighted UniFrac (Lozupone & Knight, 2005), and Bray-Curtis dissimilarity for all unique non-zero pairs of plots was the response variable (n = 861 for each metric). Standardized values for distance between continuous environmental variables were used to predict beta diversity. Grazing treatment categories were created for all possible combinations of treatments (e.g. beta diversity of grazed vs grazed, grazed vs ungrazed, and ungrazed vs ungrazed samples). We included a random effect for spatial autocorrelation that represented all unique alphabetized combinations of sites. Additionally, we used a random slope for each model.

Eleven predictor variables were analyzed for all models; grazing treatment, soil organic matter (SOM), pH, rainfall, percent sand, silt, and clay, and total concentrations of nitrogen, phosphorus, calcium, and iron. Rainfall for 2012 was as determined by satellite measurements from NASA’s Global Precipitation Measurement mission (Hou *et al.*, 2013). Means and a 95% credible interval for the posterior distribution are reported in Table 2 and Fig. 7. To estimate variation explained by each model, the ‘bayes_R2’ function in ‘rstanarm’ was used to calculate an r-squared (Stan Development Team, 2017). To determine the variation explained by the fixed effect of each model, the full model r-squared (R^2^ Full) was partitioned into an r-squared for the fixed (R^2^ Fixed) and random effect (R^2^ Random) by calculating the sum squared error for the full model and for a null model with random effects only. All models used three chains with default parameters (family = gaussian, prior = normal, iterations = 2000, adapt = 0.99), and all models converged (Rhat < 1.05).

To summarize overall patterns between bacterial community composition and abiotic factors, we used principal coordinate analysis (PCoA) and the Spearman’s rank correlation coefficient in the BIOENV function from scikit-bio. To test the effect of abiotic characteristics on bacterial community composition, we used a distance-based redundancy analysis (db-RDA) (Legendre & Anderson, 1999) performed with forward and backward model selection using the ‘capscale’ and ‘ordistep’ functions in the vegan package version 2.5-4 (Oksanen *et al.*, 2018). We performed the db-RDA analysis on Bray-Curtis dissimilarity with default parameters. Percent sand was removed from the selected db-RDA model post hoc because it did not contribute to the overall model. Indicator species analyses of OTUs (operational taxonomic units) were used to determine which bacteria were associated with grazing treatments and sites using the ‘multipatt’ function in the ‘indicspecies’ package version 1.7.6 (De Cáceres & Legendre, 2009) package in R (version 3.3.0) with default values. Only OTUs with significant indicator species (P < 0.05) are reported. P-values were determined with 999 permutations. The square root of the indicator value (func = “IndVal.g”) was used as the test statistic.

## Results

Soil nutrient analyses indicated significant edaphic gradients created by the Ngorongoro Volcanic Highlands (Fig. 1; Table 1). As expected, soil throughout the southern Serengeti plains is enriched by volcanic deposits rich in phosphorus, iron, and calcium, with deposits gradually decreasing with increasing latitude. Soil phosphorus concentration was significantly higher in the southern site (6.1 mg cm^−3^) than the northern site (0.1 mg cm^−3^; Table 1). Total soil nitrogen concentration was between 1.1 and 2.3 (mg cm^−3^) throughout the seven sites (Table 1). Soil organic matter ranged from 6.9% (BAL) to 16.2% (BRS; Table 1). Total calcium and iron concentrations were lowest in the northern site (1.8 and 4.5 mg cm^−3^, respectively) and highest in the southern site (20.2 and 24.0 mg cm^−3^, respectively; Table 1). Sites in the north were comprised of sandy loam soil with low phosphorus concentration while sites in the south were silty loam with high phosphorus concentration (Fig. 1; Fig. 2; Table 1). Because of the influence of volcanic ash, many of the environmental variables were highly correlated (Fig. S1). Notably, percent sand, silt, and clay were more than 90% correlated with each other and between 70 and 90% correlated with phosphorus concentration (Fig. S1).

**Figure 2.**
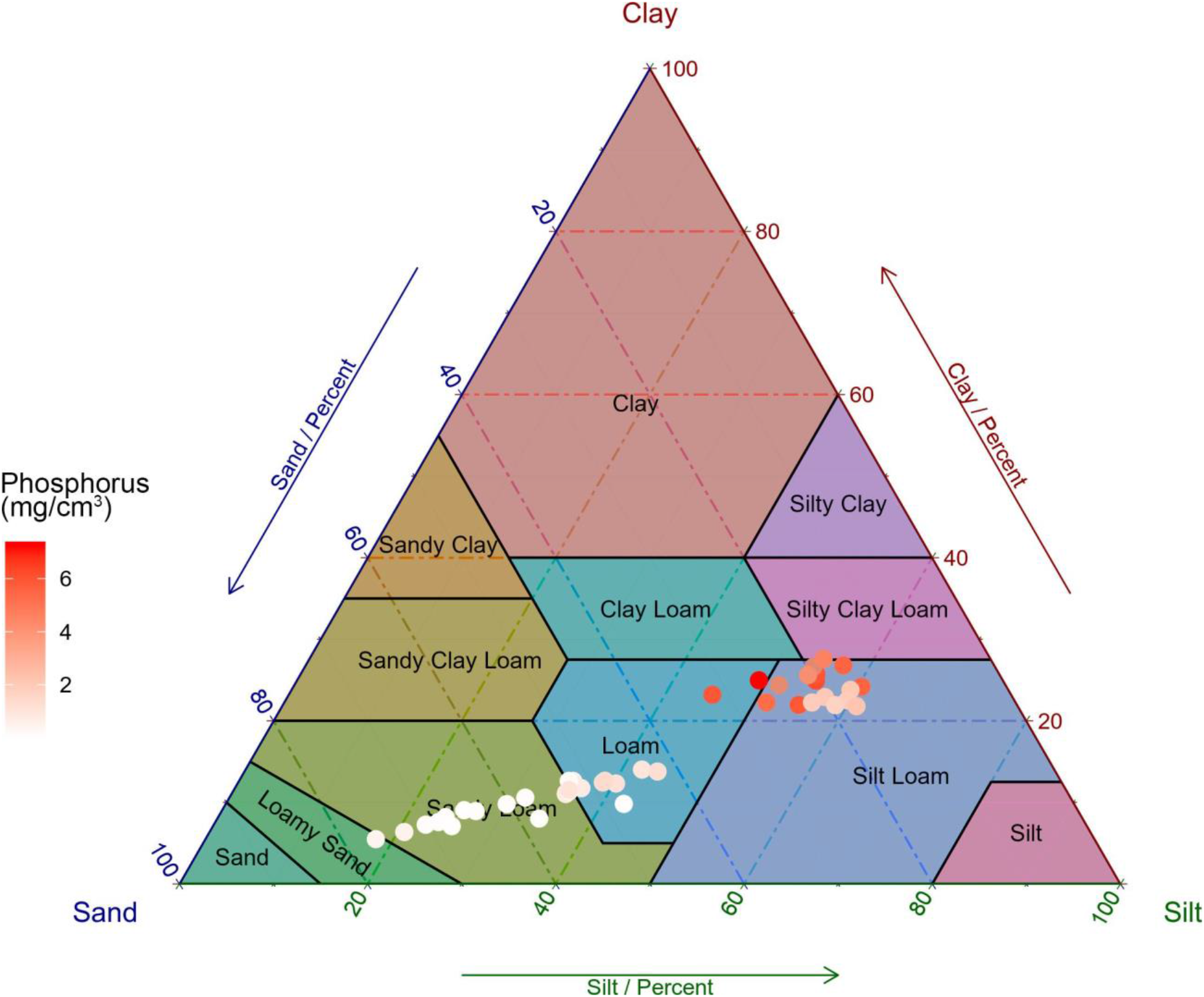
Soil texture and phosphorus concentration of each plot at seven sites in this study. Plots are colored by soil phosphorus concentration, where darker red indicates fine-textured, high phosphorus soil that result from deposition of volcanic ash from highlands south of the Serengeti National Park.

After read quality filtering, sequencing resulted in a total of 5,702,184 reads matching a total of 32,372 OTUs. To remove potentially erroneous OTUs (operational taxonomic units), we used a stringent OTU table filtering threshold that removes singletons by sample and OTUs below 0.005% of total sequence abundance (Bokulich *et al.*, 2013). OTU table quality filtering resulted in 4,431,104 (78.7%) sequences matching 2,782 OTUs (8.6%). For all 42 samples, we had an average of 105,502 sequences per sample, with a minimum of 52,139 and maximum of 162,093 sequences.

Microbial communities in this study were primarily dominated by Actinobacteria (19.5%), Proteobacteria (19.5%), and Acidobacteria (19.2%). Other significant (> 5%) phyla included Verrucomicrobia (8.0%), Firmicutes (6.0%), Chloroflexi (6.0%), and Planctomycetes (5.5%). Phyla with 5% or less relative abundance comprised 16.2% of the overall abundance. Average relative abundances for each site are reported in Fig. 3a. We used four different alpha diversity metrics to determine if environmental variables are related to richness and evenness (Fig. S2). We highlight the strong correlation between percent silt and evenness in Fig. 3c. Average Shannon diversity for each site ranged from 8.8 (± 0.2) in BAL to 9.3 (± 0.1) in BRS and was correlated positively with percent silt (R^2^ = 0.17) and total phosphorus concentration (R^2^ = 0.14). Overall, evenness was high throughout our study, ranging from 0.86 (± 0.01) to 0.91 (± 0.01). Evenness of communities was positively correlated with silt (R^2^ = 0.16), phosphorus (R^2^ = 0.25), and pH (R^2^ = 0.11), and negatively correlated with rainfall (R^2^ = 0.33). Average Faith PD, a phylogenetic diversity estimate, was lowest in BAL (61.2 ± 14.3) and highest in MSB (82.1 ± 6.0) and was not highly correlated with environmental variables. Similarly, richness as measured by the number of observed OTUs, was lowest in BAL (859 ± 242.2) and highest in MSB (1253 ± 156.2) and was not correlated with environmental variables. Relative abundances of many of the 50 most common OTUs were strongly correlated with one or more environmental variables, mainly soil texture (Fig 4; Table S2). Of the 12 strongest correlations, nine had positive and three had negative relationships with percent silt (Fig. S3; Table S2). Indicator species analysis of the grazing treatments revealed that all indicator OTUs were in the bacterial kingdom (Table S3), three OTUs were indicators of grazed plots and 31 were indicators of ungrazed plots (Figure 3b; Table S3).

**Figure 3.**
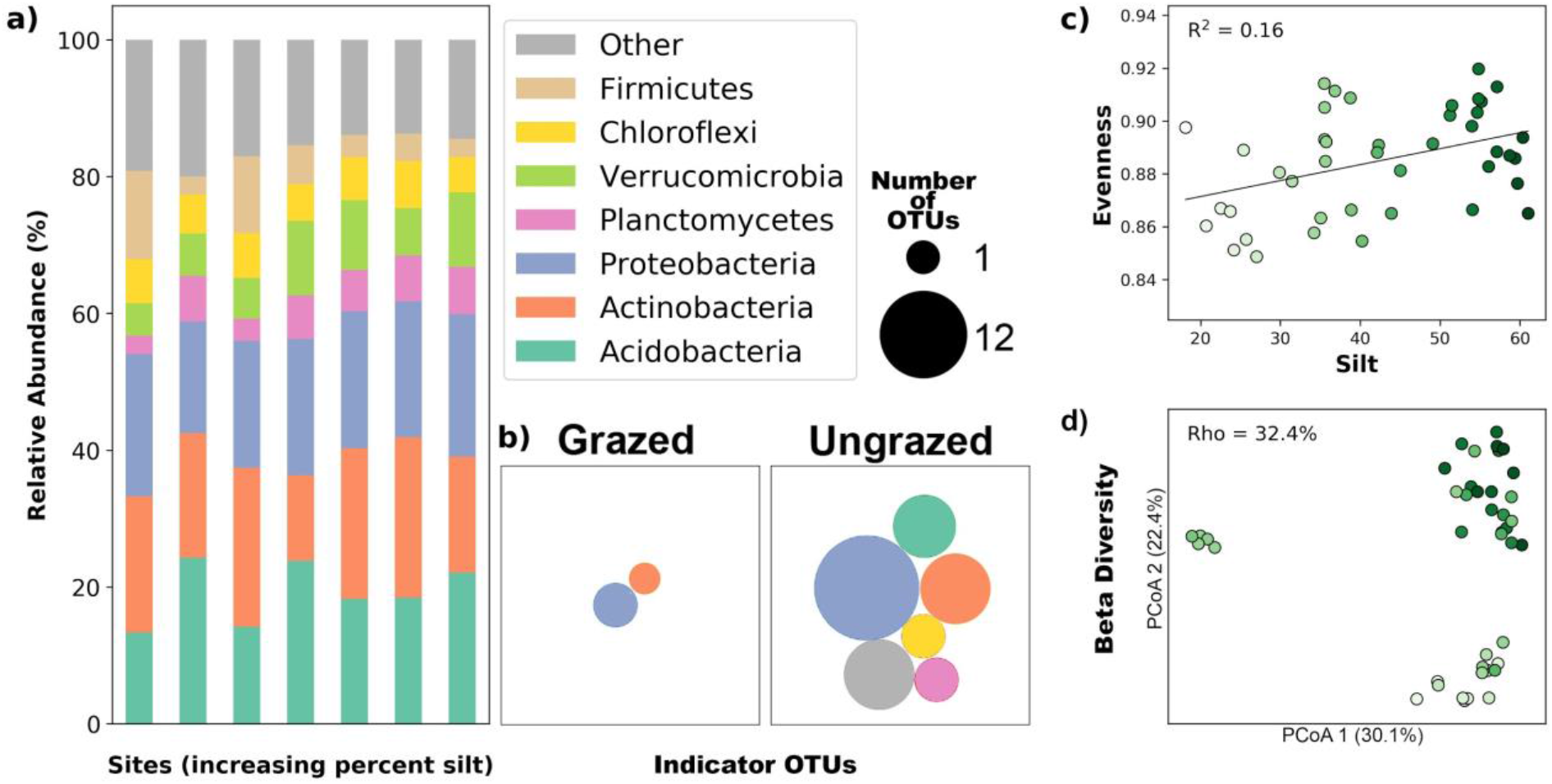
Illustrations of the bacterial community structure across environmental gradients and grazing treatments in the Serengeti. a) Correlation between Pielou’s evenness, where 1 is completely even community and 0 in uneven. Points are colored by percent silt, darker colors are higher values, to facilitate interpretation of b) beta diversity as determined by Bray-Curtis dissimilarity. Spearman’s rho value is reported to illustrate the correlation between beta diversity and percent silt (green gradient). c) Relative abundance of bacterial phyla. d) Visualization of indicator species grouped by phyla, indicated by colors. Area of each circle represents the number of indicator OTUs within each phylum.

**Figure 4.**
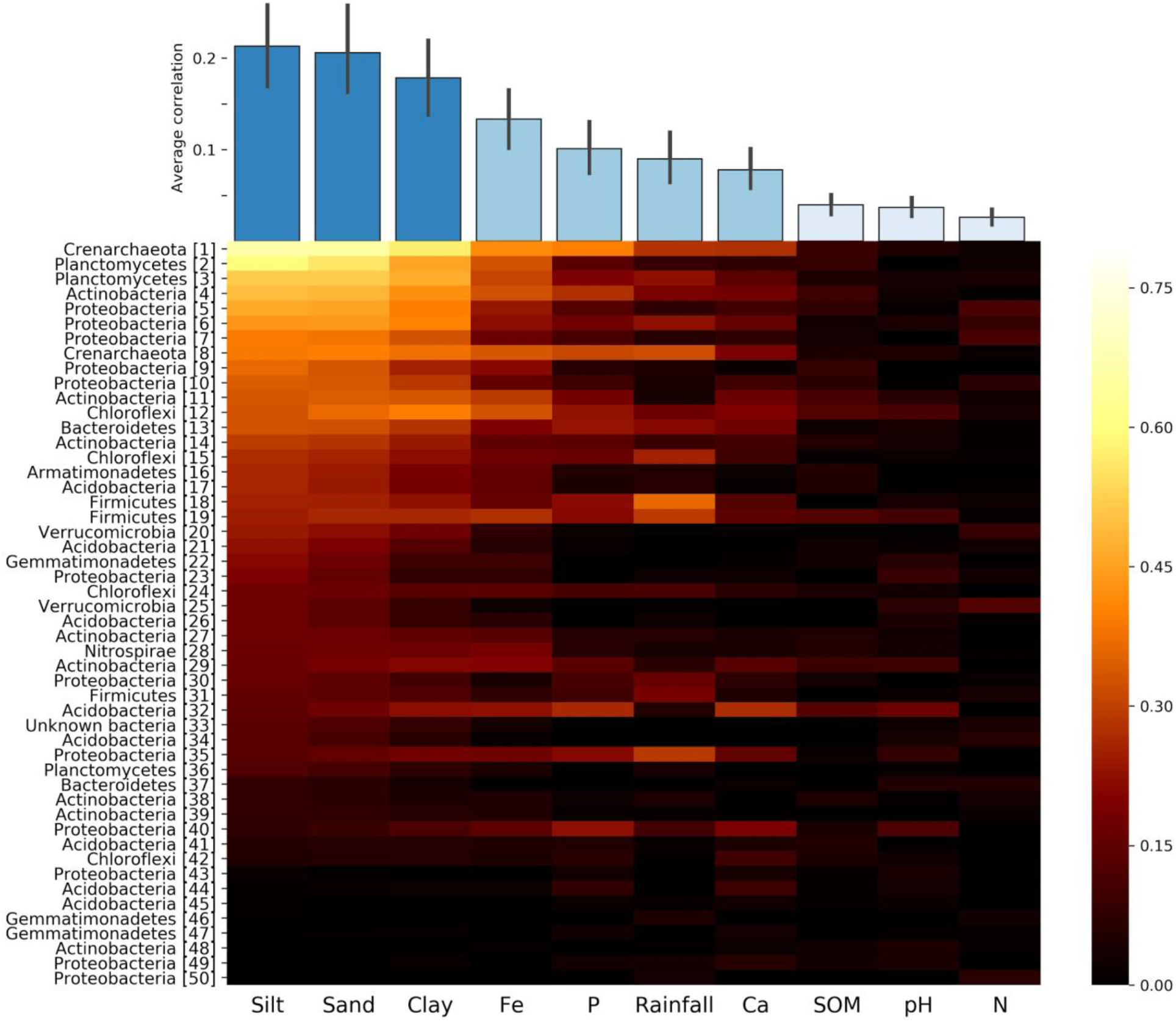
Correlation values for comparisons of relative abundance of 16S rRNA for individual operational taxonomic units (OTUs) and abiotic variables. Phyla for the 50 most abundant OTUs are displayed on the y-axis. The top-left most value is the highest correlation, and values are sorted into descending order for each abiotic variable and OTU. Blue bars above the heatmap represent the average correlation, with a 95% confidence interval, for each environmental variable. Blue shades are used to emphasize visual differences and do not represent a statistical difference.

To estimate the importance of environmental variables on the beta diversity of bacterial communities, we used spearman rank correlation coefficients with distance matrices for three different beta diversity metrics. Phylogenetic (UniFrac) and non-phylogenetic (Bray-Curtis) metrics were similar across four highlighted environmental variables, phosphorus concentration, rainfall, percent silt, and pH (Fig. S4). Despite high correlations between the measurements (Fig. S1) of phosphorus with silt (r = 0.71) and pH (r = 0.75), spearman correlations were much higher for percent silt (30.5 to 40.6%; Fig. 3d; Fig S4), and low for pH (between −7 and −2.9%; Fig. S4). Phosphorus concentration and rainfall are highly correlated with each other (r = −0.83; Fig. S1), and spearman correlations were similar for both variables and all three beta diversity metrics (Fig. S4). Much of the dissimilarity of bacterial communities can be attributed to the presence or absence of OTUs, indicated by the similarities of unweighted UniFrac correlations and Bray-Curtis (Fig. S4). Slightly higher correlations for the first two PCoA axes of weighted UniFrac, when compared to unweighted UniFrac, indicates that some variation in the bacterial communities can be explained by the phylogenetic structure of abundant OTUs (Fig. S4). In addition, we used a dbRDA to understand the effect of environmental variables on Bray Curtis distances of bacterial communities. Model selection of dbRDA indicated that soil texture, rainfall, and phosphorus and iron concentration were important in explaining variation in bacterial community composition (F = 2.6, p < 0.001; Fig. S5). Overall, the dbRDA explained 16% of the variation in community composition (adjusted R^2^ = 0.164).

To gain a deeper understanding of the relationship of the effects of grazing and abiotic characteristics on beta diversity, we used Bayesian linear mixed effects models. To avoid the multicollinearity between environmental variables, we compared single linear regressions. We used the leave-one-out information criterion to compare models. Similar to Akaike’s information criterion, lower values indicate better model fit. Model results are reported in Table S4 and visually represented in Fig. S6. Soil texture and phosphorus provided the best fit for determining the structure of bacterial communities. Results for both weighted and unweighted UniFrac were remarkably similar to those of Bray-Curtis. Percent clay, silt, and sand were three of the top four models. Total phosphorus concentration ranked third on our list of models, the 95% credible interval for phosphorus concentration in relation to beta diversity slightly overlapped zero (CI = [0.00, 0.53]), however, the mean of the posterior distribution (0.27) was the largest and was consistent with an effect, which may indicate that a large portion of the bacterial community structure is determined by levels of this important nutrient. Beta diversity values (y) range between 0.19 and 0.85, and standardized coefficients (x) ranged from 0 to approximately 3. Therefore, a mean posterior distribution for a coefficient with a value of 0.27 (phosphorus) would indicate that for every 1 unit of change in the standardized x-value, beta diversity increases by 0.27, a large difference considering the range of possible values. Linear mixed models with other environmental variables produced fixed effect correlations near zero. Model results for the grazing treatment (CI = [−0.07, −0.02]) are consistent with an effect of grazing on the beta diversity of bacteria (2% of variation explained).

## Discussion

Because the Serengeti is a relatively undisturbed grassland, we can assume that the soil microbes observed in this study represent a long-term, stable community, notwithstanding seasonal fluctuations. Furthermore, microbial populations likely reflect interactions with the macrofauna through reciprocal upward and downward trophic cascades (Sinclair *et al.*, 2010)(Anderson *et al.*, 2007; Stevens *et al.*, 2018)(Sinclair *et al.*, 2010); Stevens *et al.*, 2018). Results of this study reveals the importance of soil texture and mineral content in structuring microbial communities, and we also show that grazing by large, migratory mammals impacts microbial communities within the topsoil of the Serengeti National Park. The global distribution of bacteria has been linked to a hierarchy of correlated factors, and especially soil pH, organic matter and nitrogen availability (Fierer, 2017). Many studies have found that soil pH is a significant driver of bacterial diversity (Fierer & Jackson, 2006; Lauber *et al.*, 2009; Griffiths *et al.*, 2011; Kaiser *et al.*, 2016); but we observed pH to have little relationship with the soil microbes in the Serengeti (Figs. 4, S2-S6; Table S4). A likely reason why our results are not consistent with the literature is that the neutral soil pH range (6.3 to 7.8) observed in this study which is ideal for microbial diversity, as opposed to the larger range of pH (3 - 9) captured globally (Fierer & Jackson, 2006). Furthermore, in contrast to a global analysis showing relationships between soil microbial communities and organic matter and nitrogen availability (Fierer 2017), our study showed no relationship between microbial diversity and these soil variables (Figs. 4, S5, and S6; Table S4). Instead, we found that the richness and evenness of microbes increased in sites with finer textured soil (Figs. 3 and S2) and specific OTUs were highly correlated with texture variables (Fig. 4).

For organisms living at the scale of soil particles, the size and manner in which those soil particles coalesce can have a profound influence. Many studies have reported a connection between soil particle size and microbial diversity (Sessitsch *et al.*, 2001; Carson *et al.*, 2010; Chau *et al.*, 2011; Neumann *et al.*, 2013). Sand has larger pores with higher connectivity and therefore retains less water and nutrients than finer textured soil. The strong relationship between microbial community structure and soil texture could be linked to water potential, soil moisture, pore connectivity, and nutrient diffusion, rather than the texture itself (Saxton & Rawls, 2006; Dechesne *et al.*, 2008; Carson *et al.*, 2010; Serna‐Chavez *et al.*, 2013; Neumann *et al.*, 2013). A previous study found microbial abundance to be positively correlated with water holding capacity and soil moisture in finer textured soil in the Serengeti (Ruess & Seagle, 1994). Biotic interactions are influenced by water content, pore connectivity, and nutrient diffusion and could also influence microbial abundance and diversity. Small soil pores in fine textured soils may provide microbes a refuge from predation by bacterivorous protozoa and nematodes; and thus, loss through predation is likely to be higher in wetter, coarser soils that enable motility (Hassink *et al.*, 1993; Nielsen *et al.*, 2014). Furthermore, highly connected water-filled pores may favor competitive interactions (Treves *et al.*, 2003). Models of two bacterial species indicate less coexistence in wet soil conditions because connectivity facilitates interactions among microbes such that highly competitive species can more easily exclude poor competitors in saturated soil compared to dry soil (Hardin, 1960; Dechesne *et al.*, 2008; Long & Or, 2009). Overall, the structure of microbial communities in the Serengeti likely reflect both the top-down biotic influences of predation and bottom-up abiotic factors.

Our results indicate that environmental gradients resulting from volcanic inputs of ash influence the biogeography of microbes in the Serengeti, but it is impossible to uncouple the interconnected soil factors that arise from the volcanic deposits. Phosphorus concentration explained 14% of the variation in community composition and was consistent with a strong effect on beta diversity (Table S4; Fig. S6). Soil texture, on the other hand, explained 17% of the variation but was inconsistent with an effect on beta diversity (Table S4; Fig. S6), even with a strong gradient in soil texture (from 25% to almost 60% silt). Therefore, it is likely that both phosphorus concentration and soil texture directly and indirectly (through biotic interactions) influence microbial community structure. Results from our mixed models indicate that further research is necessary to completely disentangle the effects of soil texture and phosphorus concentration. An experimental design created to separate mineralogy from soil texture could help separate these effects.

Life in Serengeti soil evolved to co-exist beneath one of the largest mammalian migrations on Earth. We would expect the exclusion of large herbivores should have an effect on the microbial communities and our linear mixed model results are consistent with that prediction (Fig. 3b; Table S4; Fig. S6). Further, we found 34 total indicators of grazing, including one OTU within Rhizobiaceae (Table S3), some of which are known to fix nitrogen within plant roots (Spaink *et al.*, 2012). An association between Rhizobiaceae and plants within grazed plots could indicate a cooperative strategy to compensate for aboveground herbivory (Ramula *et al.*, 2019). Specifically, we found three indicators of grazing and 31 indicators of ungrazed plots (Fig. 3b, Table S3). There are many potential mechanisms by which herbivory might influence soil microbial communities (Bardgett *et al.*, 1998). The influx of nutrients from mammalian waste products could cause a shift in bacterial communities, increasing community differences between grazed plots, especially at larger spatial distances, while ungrazed plots maintain higher similarity, as our data implies (Figs. 3 and S6; Tables S3 and S4). Herbivory has altered the composition of plant communities at our sites, which in turn could alter bacterial communities (Anderson *et al.*, 2007; Prober *et al.*, 2015). Arbuscular mycorrhizal fungi can influence the community structure of soil bacteria (Artursson *et al.*, 2006), and the abundance of these fungi has been shown to be higher inside the fences that exclude herbivores and also in the southern sites with fine textured soil and higher phosphorus content (Antoninka *et al.*, 2015; Stevens *et al.*, 2018, 2020). Millions of migratory mammals offer soil microbes dispersal opportunities resulting in more homogenous communities (Vos *et al.*, 2013). Future research is needed to link the mammalian microbiome with the soil microbiome using a time-series of sampling that coordinates with the grazing cycle in the Serengeti.

Much of the biogeography of microbial communities remains a mystery, but our data provides some insights into the distribution of soil microbes in a naturally grazed grassland. This study highlights the importance of soil properties, and especially texture in structuring a significant portion of microbial evenness and beta diversity. Additionally, we discovered that the removal of mammalian herbivores had a measurable effect on the beta diversity of microbial communities. Together, these results may help inform predictions of the regional biogeography of bacteria in natural, tropical grassland ecosystems. Future studies will benefit from a deeper understanding of microbial functional diversity and the spatial and temporal dynamics of life in the Serengeti soil.

## Acknowledgements

The authors would like to thank Michael Anderson, Mark Ritchie, Jeff Propster, Anita Antoninka, Dan Revillini for their advice and input. We also thank Gustavo Zimmer for help with DNA extractions, and Lela Andrews and the NAU Environmental Genetics and Genomics Laboratory (EnGGen), Northern Arizona University, Flagstaff, AZ (nau.edu/enggen) for help with sequencing. Thanks to Emery Cowan for her contribution to the efficacy of the science communication.

## Author contributions

NCJ planned, designed the research, and collected samples. BMS performed bioinformatics, statistical analyses, and created figures. DLS and BMS created and analyzed linear models. BMS analyzed and, with the help of NCJ, interpreted data. BMS created figures and wrote the first draft of the manuscript. NCJ edited the final version.

## Figures and Tables

Table 1. Annual precipitation (AP), and edaphic properties of the seven study sites in the Serengeti National Park, Tanzania. See attached .xlsx spreadsheet

